# Automated Analysis of Neuronal Morphology through an Unsupervised Classification Model of Neurites

**DOI:** 10.1101/2022.03.01.482454

**Authors:** Amin Zehtabian, Joachim Fuchs, Britta J. Eickholt, Helge Ewers

## Abstract

Brain function emerges from a highly complex network of specialized cells that are interlinked by billions of synapses. The synaptic connectivity between neurons is established between the elongated processes of their axons and dendrites or, together, neurites. To establish these billions of often far-reaching connections, cellular neurites have to grow in highly specialized, cell-type dependent patterns covering often mm distances and connecting with thousands of other neurons. The outgrowth and branching of neurites are tightly controlled during development and are a commonly used functional readout of imaging in the neurosciences. Manual analysis of neuronal morphology from microscopy images, however, is very time intensive and error prone. Especially fully automated segmentation and classification of all neurites remain unavailable in open-source software. Here we present a standalone, GUI-based software for batch-quantification of neuronal morphology in fluorescence micrographs with minimal requirements for user interaction. Neurons are segmented using a Hessian-based algorithm to detect thin neurite structures combined with intensity- and shape-based detection of the cell body. To measure the number of branches in a neuron accurately, rather than just determining branch points, neurites are classified into axon, dendrites and their branches of increasing order by their length using a geodesic distance transform of the cell skeleton. The software was benchmarked against a large, published dataset and reproduced the phenotype observed after manual annotation before. Our tool promises greatly accelerated and improved morphometric studies of neuronal morphology by allowing for consistent and automated analysis of large datasets.

## Introduction

Neuronal function manifests most obviously their complex morphology, and functional circuits require a tight regulation of the generation of each process and branching point. Dendrites serve as input structures that perform different integrations of the axonal input. Axons, on the other hand, distribute their signals to multiple cells that may be located in multiple brain regions. The connectivity patterns of neurons are both predefined in development as well as guided by local cues during neurite extension. Altered connectivity patterns in the brain seem to underly many neurodevelopmental disorders such as autism spectrum disorder, mental retardation or schizophrenia (Calhoun et al., 2012).

The complex morphology of neurons is established during development and starts to emerge during migration of newborn neurons in the developing brain. Neuronal polarization, axon elongation, dendritic arborization and synapse formation also happen in culture and the molecular players are largely conserved, making cultured hippocampal neurons an established model for the development of a functional neuronal morphology (Dotti et al., 1988; Li and Sheng, 2003; Polleux and Snider, 2010; Cembrowski and Spruston, 2019; Denoth-Lippuner and Jessberger, 2021).

The broad variety of neuronal cell types shows a wide diversity of morphological parameters that are related to function. Neuronal cell types can vary in number, length and branching of axons and complexity of dendritic arbor. The quantification of neuronal morphology via extraction of basic parameters such as neurite length, number of branches, and degree and density of branching points from microscopy images is thus of fundamental importance in the neurosciences. One of the longest used assays is Sholl analysis (Sholl, 1953), which approximates dendritic branching by counting the number of times neurites cross concentric circles emanating from the soma at increasing distances. Sholl analysis is a regular feature of neuronal segmentation software. The commonly implemented strategies range from manual tracing over semi-automatic detection (Schmitz et al., 2011) to segmentation-free analysis (Ferreira et al., 2014).

Due to their tortuous and elongated growth patterns, however, Sholl analysis is not suitable to quantify axon morphology. Axonal outgrowth has been mainly quantified by the length of the longest process, the summed length of all axonal processes, and axon complexity as the number of branch points per micron (Sainath et al., 2017; Pan et al., 2019). Extraction of these parameters require first the reconstruction of the neuronal outline and secondly the subsequent classification of processes as axons and dendrites and their primary, secondary, and tertiary branches. Commonly implemented strategies mainly use manual tracing and classifications using tools like NeuronJ (Meijering et al., 2004).

Automated reconstruction of neurons has made significant progress over the past years, also stimulated by an interest in connectomics in projects such as the BRAIN Initiative (http://www.braininitiative.nih.gov/) or the Human Brain Project (http://www.humanbrainproject.eu/), the DIADEM challenge (http://diademchallenge.org/) or the BigNeuron project (reviewed in Parekh and Ascoli, 2013; Magliaro et al., 2019). Most current pipelines employ pre-processing strategies such as denoising and deconvolution followed by segmentation algorithms differing by the strategy they employ to distinguish neurons from background. These pipelines are implemented in commercial (Neurolucida, IMARIS, Amira, HCA-Vision) as well as free standalone packages (Neuronstudio: Rodriguez et al., 2008; Neutube: Feng et al., 2015) or as plugins in image processing software such as Matlab (TREES toolbox: Cuntz et al., 2010), ImageJ (PhD-filtering: Radojević and Meijering, 2017) or Vaa3D (APP2: Xiao and Peng, 2013; Rivulet: Liu et al., 2016; Ensemble neuron tracer: Wang et al., 2017; NeuroGPS-Tree: Quan et al., 2016; Advantra: Radojević and Meijering, 2019). An enormous number of further solutions have been also developed over the years to cope with segmentation and reconstruction of the neuron morphology, resulting in a vast number of different applications (Narro et al., 2007; Oberlaender et al., 2007; Schmitz et al., 2011; Meijering, 2010; Peng et al., 2010; Peng et al., 2014; Megjhani et al., 2015; Acciai et al., 2016; Magliaro et al., 2017; Yoon et al., 2017; Ikeno et al., 2018; Abdellah et al., 2018; Shahbazi et al., 2018; Wang et al., 2019; Abdolhoseini et al., 2019; Vidotto et al., 2019; López-Cabrera et al., 2020; Bates et al., 2020).

While these works mainly focus on the reconstruction of neurons, a reliable quantification of neuronal morphology requires the subsequent classification of neurites. Recent work describes the extraction of growth parameters (Narro et al., 2007; Scorcioni et al., 2008; Billeci et al., 2013), the modelling of these parameters to derive growth or electrophysiological characteristics (Cuntz et al., 2010; Ascoli and Krichmar, 2000) and the classification of cell types based on morphology (Armañanzas and Ascoli, 2015). These advances, however, so far do not allow for automated analysis, especially of axons, which would greatly enhance throughput in image analysis and create compatibility of this assay with high-throughput approaches.

To overcome this bottleneck and accelerate data analysis, we here developed a software that allows for batch processing of raw fluorescence micrographs of cultured neuronal cells to extract morphological parameters of axons and dendrites. We set out to use concepts and tools from the fields of automatic reconstruction and unsupervised image classification to facilitate the analysis of molecular mechanisms underlying the growth patterns of primary neurons in 2D culture. Our goal was to implement the quantification of intuitive biological descriptors in a tool capable of batch processing microscopic images of neurons with a focus on developing axons where automated analysis strategies are sparse. Here we present an open-source MATLAB-based software capable of classifying individual processes of neurons as dendrites or axons and their branches, respectively in an automated manner and benchmark it against human classification and data analysis.

Our software imports raw fluorescence micrographs of neurons from standard image file formats and applies a set of consecutive tools for denoising, segmentation, detection of the soma, primary neurites and finer, second and third order branches. This is followed by unsupervised classification of the neurites. Finally, quantitative measurements are exported for each batch as a comprehensive collection of figures and tables. The exported results include, but are not limited to the size of the soma, the total length of the axon, number and the respective lengths of the axonal branches, dendrites and dendritic branches. This information can be accessible for more specific downstream processing (Narro et al., 2007; Zhou et al., 2013; Gillette et al., 2015; Acciai et al., 2016; Mihaljević et al., 2018; Abdellah et al., 2018).

We demonstrate the capability of our software on fluorescence micrographs of primary hippocampal neurons and compare the achieved neurite classification with results obtained by manual image analysis as ground truth. We find that the analysis of our software matches the analysis of an experienced user while being capable of simultaneous analysis of tens to hundreds of images in a short time without manual intervention.

## Results

### The GUI

Here we developed a graphical user interface (GUI)-based open-source software to analyze fluorescence micrographs of neurons for quantitative analysis of neurite length and branching. Files are automatically imported, and visual and numeric results are saved as standard file formats for further downstream processing and analysis. The main interface is shown in **Figure 1**. The workflow contains modules for importing data, pre-processing, segmentation, neurite classification and the export of quantitative measurements (see also **Fig. S1)**.

**Figure 1:**
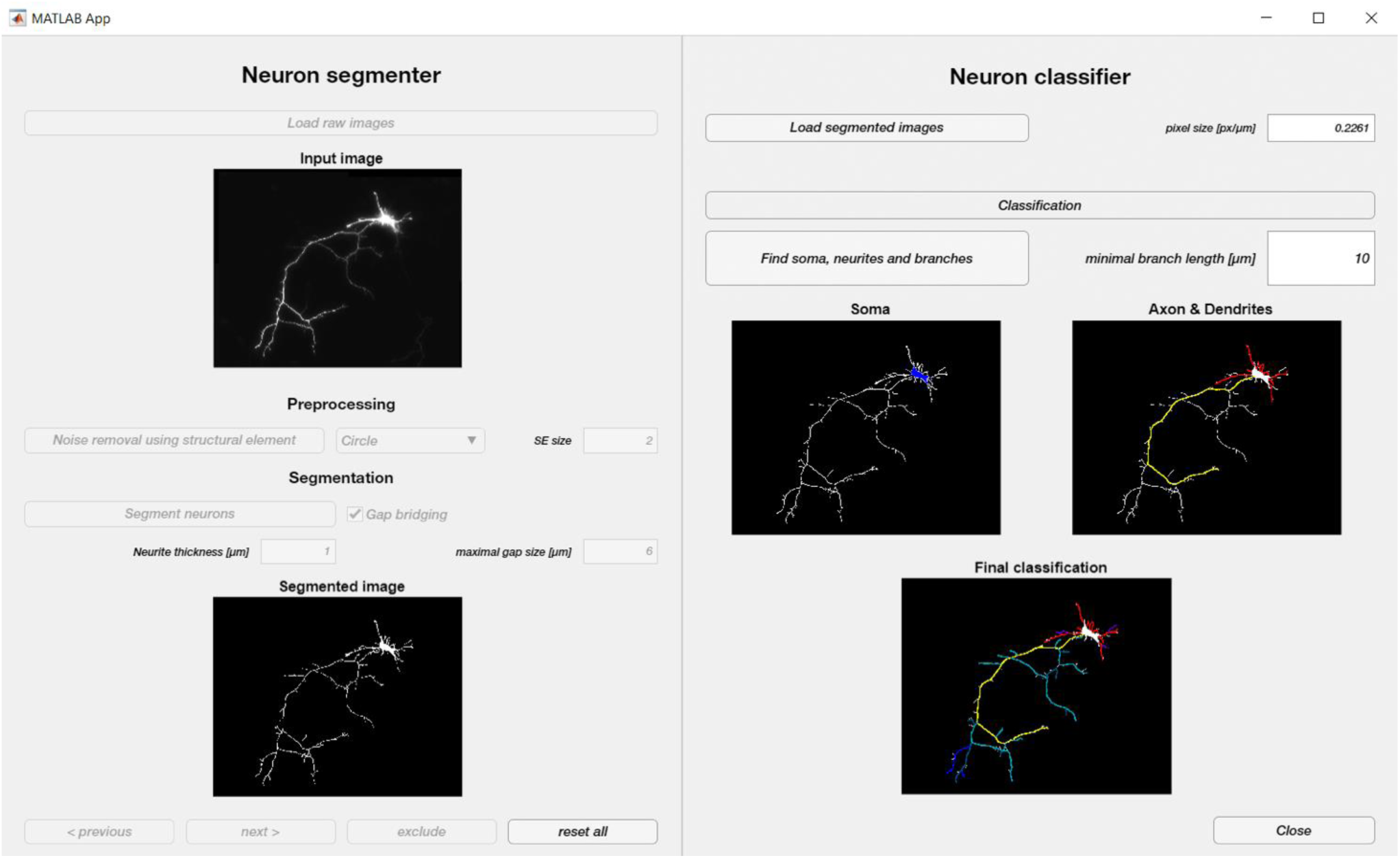
Graphical user interface and processing steps. Screenshot of the automated neuron reconstruction and neurite classification software’s GUI. Loaded raw or preprocessed images are shown as preview (left top) and can be preprocessed. The segmentation can be performed in automated or user-guided manner (left bottom). The segmented neuron is then analyzed (right), the soma is detected (blue) and its neurites are classified (axon yellow, primary axon branches teal, secondary axonal branches blue, dendrites red) and quantified.

The first block within the proposed framework allows the user to load the input data in two ways: (1) loading raw neuronal images, or (2) importing data which have been segmented or skeletonized before. The software can operate in two different modes. In default mode, the user can change parameter settings (such as pixel size, minimal soma size, minimal neurite length) for each image individually.

In batch mode, the program sequentially processes all imported data with the same settings. When the batch mode is enabled, it is essential that pixel size metainformation is included in the image file.

After loading images, an optional pre-processing is available to smoothen the image based on two different types of morphological filtering (Zehtabian and Ghassemian, 2016). Smoothing the data often results in higher quality segmentation, ensuring smoother outlines of the segmented neurons and less discontinuities.

The data will be then fed into the built-in segmentation module which follows an automated, fast algorithm based on Hessian filtering. The segmented images will then be skeletonized using standard techniques. If previously segmented data are loaded, they will directly proceed to the classification step. The segmented neurons are then processed to automatically extract the soma, axon, dendrites, axonal and dendritic branches, respectively. Quantitative measures such as the region occupied by the soma, total length of the neurites as well as the length of each neurite per se will be then extracted from the neurons. All numeric results and variables of interest are saved in a Matlab file and as .txt for further statistical analysis. A labelled .tiff image is also generated and saved for inspection of classification results.

### Segmentation

One of the critical steps in morphological analysis is segmentation, i.e., the determination what is part of the neurite and what is not. This can be challenging for fluorescence micrographs of neurons where local differences in intensity gradients can lead to artifacts such as continuity breaks or complete loss of branches. We addressed these concerns in two ways. First, we pre-processed the data to generate a homogenously high quality of input images as described before, and secondly, we equipped our segmentation module with multiscale filters that emphasize features in images that are important for efficient neurite reconstruction. A schematic overview of the proposed neuron segmentation method is depicted in **Fig. 2A** while example results of applying each step to a given neuron is shown in **Fig. 2B**.

**Figure 2:**
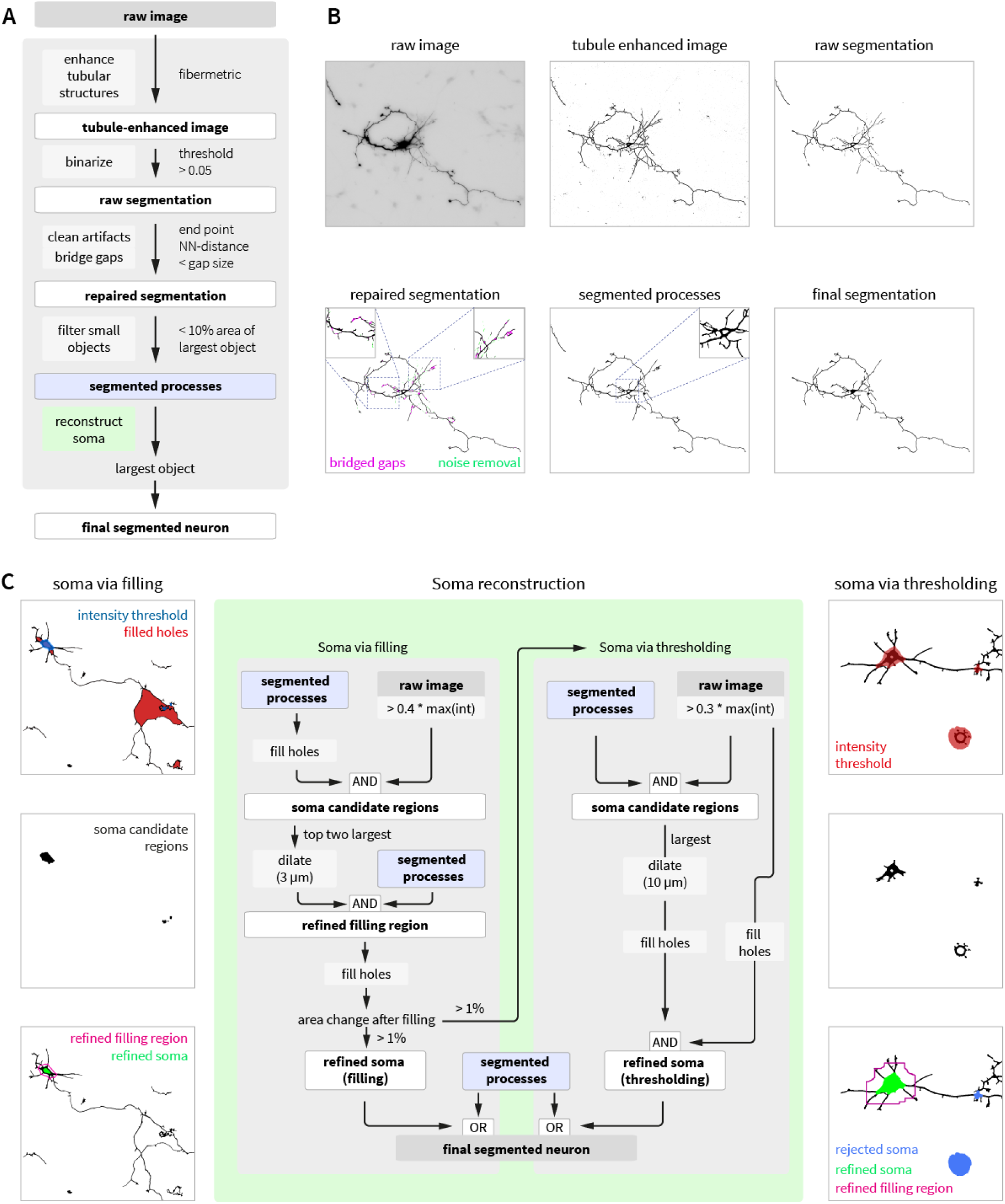
Schematic overview of the algorithm used in the built-in segmentation module. **(A)** General overview of the data processing pipeline of the proposed segmentation method. **(B)** Visualization of results from the individual segmentation steps performed by the proposed technique with an emphasis on quality control measures that “repair” errors in segmentation (pink and green, bottom left). **(C)** The soma reconstruction scheme: morphological filling serves as the preferred reconstruction method but is backed up by intensity thresholding if unsuccessful.

The first step of the proposed segmentation algorithm highlights elongated linear structures (which are the neurites in our case). To this end, we employ a technique proposed for the detection of blood vessels (Frangi et al., 1998), which is based on eigenvalue analysis of the Hessian. Hessian-based multiscale filtering extracts information related to local second order structures within the image. This *fibermetric* algorithm enhances linear structures of a specified thickness, while minimizing information on thinner or thicker structures. In our software, the starting parameter for this ‘*neurite thickness*’ (in µm) may be adjusted to accelerate performance of the software if prior knowledge exists. Otherwise, the software will adaptively set the value of this parameter.

To create the neuron model, we apply an intensity-based hard-thresholding on the tubule-enhanced image to obtain a binarized image. Hard thresholding potentially introduces short gaps within the neurites. This may be caused by low contrast in images or locally varying thickness of neurites, often resulting in incomplete skeletonization. To overcome this problem, the software includes an optional gap-filling algorithm which is based on finding and connecting nearest neighbors of endpoints from unconnected objects. If the closest endpoint is closer than a maximal allowed user-specified gap size (in µm), the gap is closed with a direct line. Bridging gaps can successfully re-connect short gaps between neurites (**Fig. 2B**, repaired segmentation left inset). However, it should be used with caution, as also noise or neighboring short processes might be artificially hyperconnected when large gap sizes are allowed for this algorithm (**Fig. 2B**, repaired segmentation right inset).

As can be inferred from **Fig. 2B** (segmented processes), although the linear structures are segmented well, the interior region of the soma is not segmented. To accurately reconstruct the soma, we apply morphological operations to the binary image to fill interior image regions and holes (**Fig. 2C**). However, individual neurites may cross paths and thus create closed-off interior regions other than the soma (**Fig. 2C**, top left). To overcome this problem, our algorithm screens for soma candidate regions (**Fig. 2C**, middle left) with a high average brightness. The largest two of such overlapping regions are subsequently dilated and serve as region selectors for the filling operation. (**Fig. 2C**, bottom left). In this way morphological filling is only applied around the most likely position of the soma to avoid mis-localization of the soma or overestimation of soma size.

While this approach works well for the majority of tested neurons, in some cases the soma region is not fully enclosed after segmentation and thereby cannot be morphologically filled. Such cases are detected and soma regions are then reconstructed using an intensity threshold only. To limit the effect of brightness artifacts on the image, the algorithm first finds new soma candidate regions based on the amount of overlap of the intensity threshold and the segmented neurite skeleton and then only reconstructs the soma with largest overlap to the neuron (**Fig 2C**, right panels). To avoid contribution of small segmentation artifacts and noise to this overlap quantification, unconnected small objects are removed prior to soma reconstruction. The final step of segmentation finds the largest connected object on the image and removes all unconnected objects, thereby focusing the analysis to only one cell.

Our software also allows for use of the automated classification only and is compatible with data segmented manually in Vaa3D (Peng et al., 2010) or SynD (Schmitz et al., 2011). It is possible to load segmentations from those tools as binary image files and then to proceeding with the subsequent classification step in our software only.

### Classification

The neurite classification scheme is illustrated in **Fig. 3**. The classification step initiates with the detection of the soma, followed by a stepwise detection of the longest processes originating from the detected soma. The longest of these processes is classified as the “axon”, the remaining processes as “dendrites”. Classification proceeds by detecting processes originating from the axon as “primary branches” and subsequently processes originating from these primary branches. Detection is completed after no further level of branches is detected or a hard cap of maximal 10 levels of branches is reached. After detection of axonal branches, the same procedure is applied to all dendrites and their branches (**Fig. 3**).

**Figure 3:**
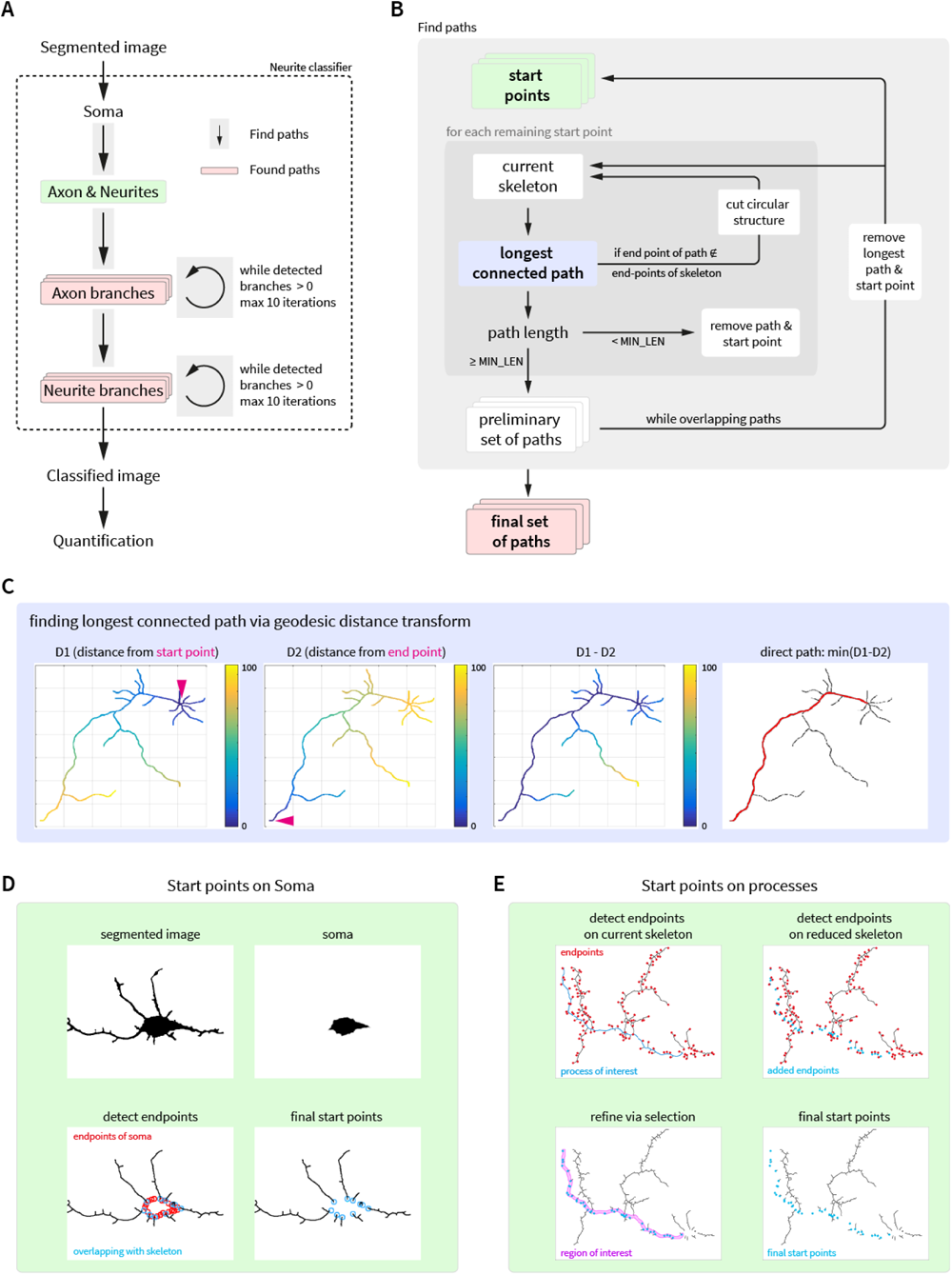
Illustration of neurite classification. **(A)** General overview of the proposed classification approach. Axon, dendrites and their branches are classified using the same algorithm ‘find paths’. Axon branches are classified iteratively as primary branches, secondary, etc. before dendrite branches. **(B)** Strategy for ‘find paths’ used to detect the set of non-overlapping longest paths from a set of starting points. For each start point, the longest direct path in the skeleton is determined using a geodesic distance transform. In case the skeleton contains circular features precluding length measurements, these structures are separated at the originating branch point. In case a detected path does not exceed a user-specified minimal length, it is removed from further analyses. In case paths from multiple start points overlap, the longest path is stored, and all other start points are reevaluated on a skeleton without this path. **(C)** Determination of the longest direct connection of two points by finding the minimum of the difference of the geodesic distance transform for both points. **(D-E)** The reliable detection of start points defines the reliability of the classification algorithm, as paths without start points will not be classified. **(D)** Start points on the soma are detected as morphological end points of the soma that overlap with the skeleton excluding the soma. **(E)** Start points on neurites (i.e., start points for branches) are detected as the added morphological end points of the skeleton after removing the given neurite (or set of neurites) from the skeleton. To refine the selection, only points within a 5 px distance (‘region of interest’) to the given neurite are retained.

### Soma Detection

Since axonal and dendritic trees both emerge from the soma, their location is required as a starting point for reconstruction (Meijering, 2010). In general, even an approximation of the soma contour is often difficult to be accurately reconstructed (Abdellah et al., 2018). One solution to facilitate detection of the soma is to stain for DNA in a separate image channel (Meijering, 2010) to identify the soma unambiguously. This does, however, not allow for an accurate measurement of soma size. In the presented work we assume that neuronal images are imported to the software as single-channel data of fluorophore-filled neurons.

**Fig. 4** illustrates our proposed soma detection scheme in detail. The procedure starts by detecting the largest connected component of a minimal diameter by a repeated morphological erosion operation using a square (with size *l*_*i*_ × *l*_*i*_ at the *i*^*th*^ step) as the structural element *SE*1 with varying size. The *SE*1 size follows a descending order starting from *l*_1_ = *S*^*SE*^ (which can be set either manually or automatically). Although the erosion operation itself reduces a given structure to a slightly thinner version, since the *SE*1 size decreases, each step of performing the repeating erosion block slightly expands the eroded image (which is called 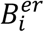 at the *i*^*th*^ step). At each step, the size of 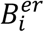 is compared to the pre-specified minimum allowed size of the soma *S*^*Soma*^. The repeating erosion cycle continues up to the point when the size of 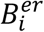 is no longer smaller than the threshold *S*^*Soma*^. At this point, the eroded image will be checked for the number of connected components (called ‘island’ hereafter). In case of having multiple islands, only the largest connected component will be kept and named as *B*^*lcc*^. The main soma detection step initiates with the centroid of *B*^*lcc*^found in the previous step. The centroid pixel (also called ‘seed point’) is considered as the initial soma (*Soma*_0_). The proposed technique then carries out a local search around the seed point and expands it by adding those pixels within its immediate spatial vicinity with intensities between *Mean*(*Soma*) ± *tol*, in which *tol* is the tolerance parameter.

**Figure 4:**
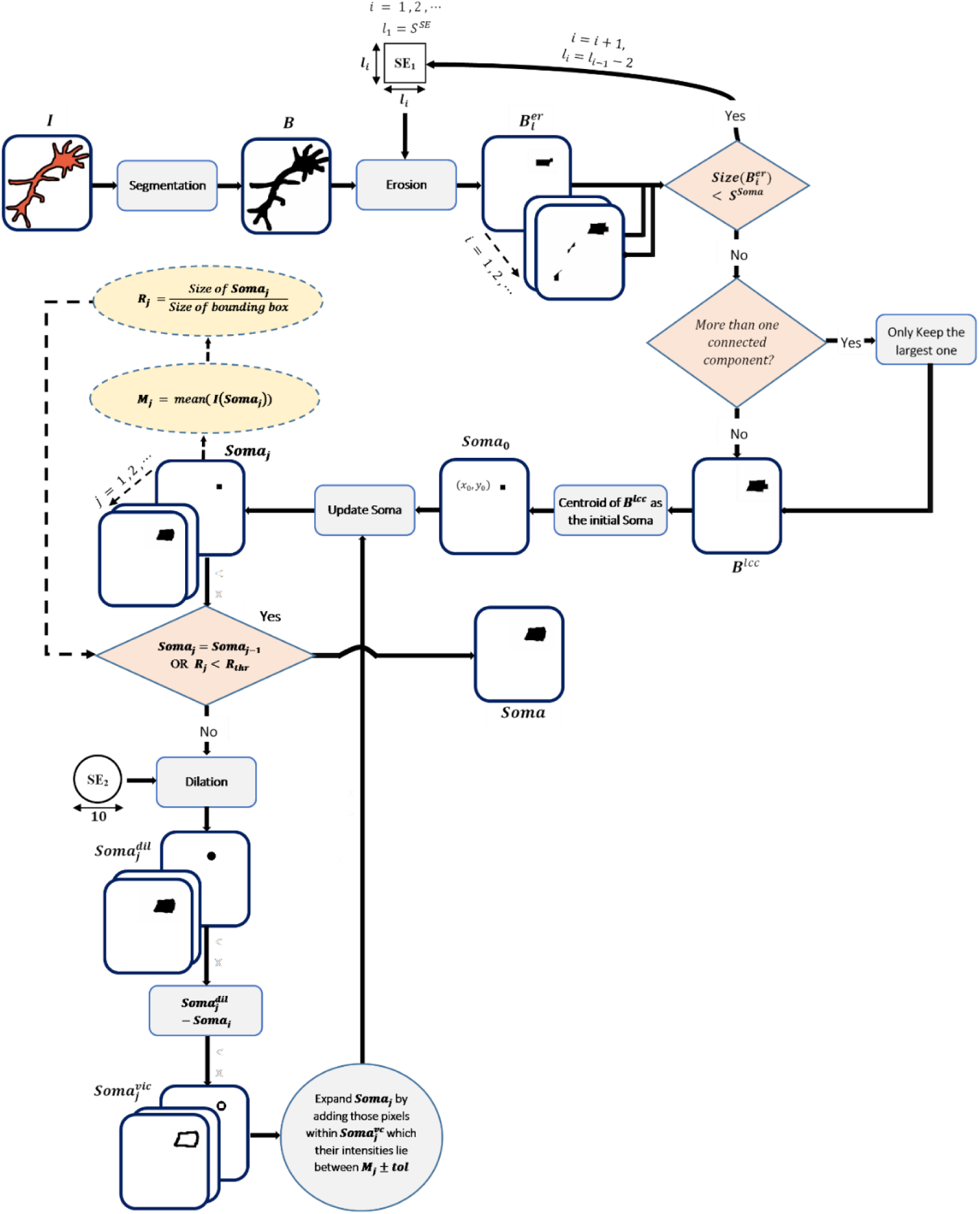
The soma detection scheme highlighting the individual steps in analysis.

The region growing step continues as long as the two following criteria are met:

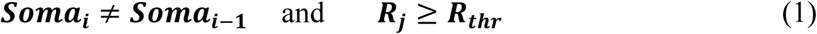

in which

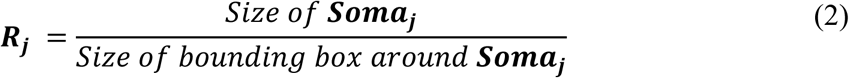

and ***R***_***thr***_ is an upper-bound threshold for the above-mentioned ratio ***R***_***j***_ with the default value set to 0.4 that can be also tuned in the Software. The parameter ***R***_***j***_ is defined to cope with ambiguities in the transitions from soma to dendrites (**Fig. 4 – Fig. S2**) .

We visually inspected the results of applying our proposed soma extraction method to several hundred neuronal images with different shapes and morphologies and found that the reconstructed soma masks delineated the real cell body in the neuron images with a high level of accuracy.

### Neurite Classification

The detection of the longest connected path originating from a specified set of start points lies at the core of the neurite classification scheme (**Fig. 3B**). We here define an axon as a single process growing individually for the longest possible distance from the soma. Beginning with soma detection, the segmented image is transformed into a skeleton using a repetitive morphological thinning operation which thins the segmented foreground to its medial axis (Lam et al., 1992). Afterwards, geodesic distance transforms (Soille, 2013) starting from specified points on the skeleton are calculated **(Fig. 3C, transform D1)**. If the maximal geodesic distance from a given start point is found on an endpoint of the skeleton, this endpoint serves as the start of a second geodesic distance transform **(Fig. 4C, transform D2)**. The difference of D1-D2 reaches a minimum at the directly connecting path of start point and detected endpoint and is stored as a detected path.

Circular structures in the skeleton complicate the detection of a longest path, as multiple paths could be equally distant to the point of maximal distance. In such a case, the maximal geodesic distance does not coincide with an endpoint of the skeleton and the detected path would be incomplete. To overcome this problem, a repair function is initiated to trace back to where the circular structure in the skeleton starts. This is achieved by walking backwards from the point of maximal distance until the respective geodesic distance is only found once in the array of distances – representing the point where two equally long paths branch out. The repair function cuts one of the paths (randomly) by removing the first pixel after the branchpoint. Subsequently, the same starting point will be used to detect the new longest geodesic distance in the skeleton until it this distance is reached at an endpoint of the skeleton.

Further steps of the path-detecting algorithms mainly deal with exclusion criteria for any specific path. If a detected path is below a user-defined minimal length of interest (the standard value is set to 10 µm as a threshold to distinguish precursor protrusions from mature branches but can be modified by users depending on the specific research question), the starting point in question and the path will be deleted from the skeleton to accelerate downstream detection. Only paths longer than this maximal length are stored for further processing.

As paths from different given starting points tend to overlap if there are intersections in a skeleton, the final step deals with removing these overlaps. If overlaps are detected, the longest path and its start point in the set are saved and subsequently removed from the analyzed skeleton, and the detection is restarted with all starting points on the modified skeleton. Restarting detection for all paths has turned out to be necessary as newly detected paths on the refined skeleton have been found to often overlap with previously detected other branches, and in rare cases, more than two branches overlapped. To avoid overlap between any branches also after this re-evaluation step, this overlap-removing algorithm is repeated until no overlap is detected between any branches. The final step of the path-detection from a set of start points removes all detected paths from the skeleton for subsequent detection of higher order branches.

The accuracy and speed of the proposed classification algorithm depends strongly on the choice of start points. For the initial step of detecting axon and dendrites, start point candidates are found as the endpoints of the detected soma. Those endpoints that overlap with the skeleton of the segmented image excluding the soma classify as final start points for axon and dendrite detection (**Fig. 4D**). Axon and dendrite detection then proceeds on the skeleton without the soma, to avoid detection of the axon-endpoint from all possible start points (the geodesic distance for unconnected parts of a skeleton is infinite and will therefore never qualify as a path of minimal distance).

To detect start points for higher order branches on axons, dendrites or their branches, endpoints and branchpoints are detected before and after removing the originating path from the skeleton (**Fig. 4E**). Additional endpoints and lost branchpoints serve as candidate start points for the detection of the next order of branches. These candidates are refined by proximity to the process of interest (within 5 px) and the refined set will initiate the first iteration of the find paths algorithm described above (**Fig. 4B**).

This selection of start points on neurites is not capable of distinguishing crossing of a neurite from branching and therefore slightly overestimates the number of branches (see next section). The implemented length threshold for branch-detection nevertheless improves accuracy of detecting relevant branches when compared to simply determining branch points as pixels with more than two neighbors: morphometric branchpoint detection does not account for the length of connected processes and is therefore more susceptible to noise in images or overly detailed segmentation results – in many cases, these paths contain only individual pixels. Furthermore, detection will speed up with iterations, as parts of the skeleton that are too short to be relevant will be removed from downstream calculations.

After extracting features of interest from the classified neurites, all numeric results as well as a labelled neuron are saved in folders specified for storing data. The numeric outputs are exported to space-delimited .txt and .csv file formats to be imported later into other software for further analysis. The segmentation and classification results are saved as .tiff files in specific folders. Additionally, all variables of interest used in the current setting will be stored in the Matlab file ‘result.mat’ as a global data matrix.

### Benchmarking

In order to evaluate the performance of our proposed framework, we compared the results of our algorithm with the analysis of an experienced user as ground truth. To do so, we used data from manual segmentation performed in a recent study on axonal branching (Brosig et al., 2019).

This dataset consists of 883 fluorescence micrographs of neurons, 204 of which were excluded from being analyzed by the software due to either the presence of large imaging artifacts, their very low signal-to-noise ratio or overlapping signal from several cells. While this means that 24% of the data are lost, this dataset has not been specifically acquired for automated analysis and a dedicated dataset would not have required such a step. If users limit their microscopy to non-overlapping cells (also non-overlapping with debris), we expect a much higher applicability. Nevertheless, we advise users to quality control segmented images before proceeding with classification to improve the quality of quantitative readouts. The remaining 679 images were fed into the proposed segmentation and classification steps (**Fig. 5A)**. In 2 images, the software produced a segmentation artifact precluding further classification (no soma reconstructed), and in 8 images the software did not label all major processes and their branches. The remaining neurons were fully segmented.

**Figure 5.**
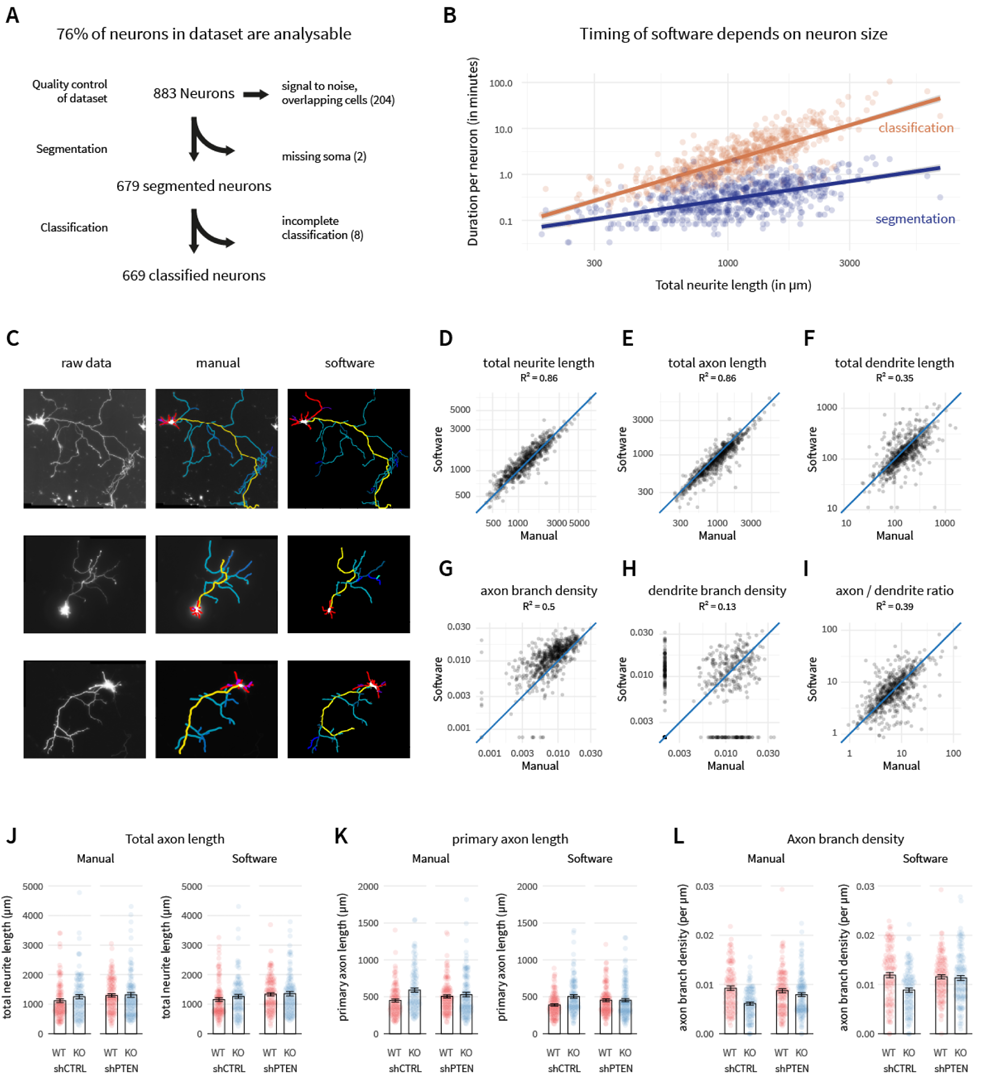
Benchmarking of software against a published dataset. (A) The test dataset consists of neurons manually analyzed for a study on axon branching (taken from Brosig et. al., 2019). (B) Duration of the calculation scales exponentially with neuron size. (C) Example results from the proposed neurite classification framework in comparison to expert user. (D-I) Quantitative comparisons of automated software analysis with expert user analysis. Axes in F-I are logarithmic due to the non-linear distribution of the data. Blue line and R-values are shown to visualize offset from unity between software and manual annotation. (J-K) Re-analysis of an experiment from Brosig et. al., 2019 (axon arborization in PRG2/PLPPR3-KO neurons with or without additional knockdown of PTEN). While absolute values are not identical, differences between biological treatments are reliably detected and not skewed by the software.

The analyses presented were run on a laptop (Intel® Core™ i7-8565U CPU 1.80GHz, 8, 16 GB RAM, Windows 10 64bit) using Matlab R2021a. Neuronal images used in our experiments are between 1.3 MB (1280 × 1024 pixels) to 29.6 MB (6720 × 4506 pixels) in size and in 8-bit depth. The median analysis time per neuron was at 2 minutes for this dataset. As shown in **Fig. 5B**, however, the required processing time scales exponentially with image properties such as image size and neuron complexity. The neurite classification is the most time-consuming step ranging between few seconds and an upper border of two hours for one image. In case the desired dataset consists of larger neurons, computation time could therefore increase considerably. However, our experiments show that even for large, highly complex images, dozens of neurons can be batch-processed over-night in a fully-automatic manner with no need for manual interventions.

Example results are presented in (**Fig. 5C)** showing the performance of the proposed method compared to the manual segmentation and classification of neurites. In many cases, the software reproduces the manual analysis nearly identically (**Fig. 5C**, top). In some cases, the expert user included additional information such as neurite intensity or crossing of neurites to define the full path of an axon differently (**Fig. 5C**, middle). In other cases, the software provided a more accurate length measurement than human users and therefore chose a different endpoint for the axon (**Fig. 5C** bottom). In general, however, reconstruction and classification of neurites appeared largely similar to expert user analysis.

To better evaluate the accuracy of the measurements generated by our software, we compared quantitative readouts of morphology of the software to those achieved by manual analysis made by the expert user (**Fig. 5D-I**): Total lengths of all reconstructed neurites, axon and dendrites serve as a measure for neuron size, densities of axonal and dendritic branches as a measure of complexity, and the ratio of axon to dendrite length as a readout for neuron polarization and therefore correct classification of the axon.

The comparison of total length measurements reveals high levels of correlation between sizes of neurons, and their axons and dendrites detected by the software and those manually reconstructed by the expert (**Fig. 5D-F**). The neurons in this dataset were imaged at a timepoint where dendrites are not fully developed and therefore still very short. Therefore, the dendrite length (**Fig. 5F**) as well as the ratio of axon to dendrite length (**Fig. 5I)** vary by orders of magnitude and are presented on a logarithmic scale. Nevertheless, in both readouts automated analysis correlated well with manual quantification, indicating the correct automated detection of the axon. The current version of this software classifies crossing points of neurites as branches, and therefore slightly overestimates branch densities (**Fig. 5G-H)**. The axon branch density nevertheless correlates well between automated and manual analysis and therefore provides a good estimate for neuron complexity. The dendrite branch density is again difficult to interpret due to the short length of dendrites in this dataset with most neurons not containing any branches on dendrites.

Despite the slight overestimation of neuron complexity, this software nevertheless is sensitive and accurate enough to detect the same phenotypes between biological treatments as an expert user (**Fig. 5J-K**). The analyzed dataset has been used to establish that PRG-2/PLPPR3 knockout does not alter total axon length (**Fig. 5J**, manual), but increases the primary axon length (**Fig. 5K**, manual) by exhibiting less axonal branches (**Fig. 5L**, manual), with additional knockdown of the phosphatase PTEN (shPTEN) rescuing this phenotype (compare to Fig 7C in Brosig et. al., 2019). The automated analysis of the same neurons provides the same differences between treatments even though absolute values of primary axon length are slightly lower and axon branch density are slightly higher compared to the manual dataset. Given the fully automated implementation, we therefore believe this tool allows for the faster screening of multiple biological treatments and for analyzing more neurons to achieve better statistical confidence.

## Discussion

With the introduction of large field of view cameras and automated microscopes producing large amounts of data in a short amount of time, the automated computational processing and analysis of morphometric data has become more important for neuroscience research. At the same time, automation offers great prospect in terms of data reproducibility, as experiments can be performed with many datapoints and stronger statistics, allowing for the detection of smaller effects. Furthermore, manual processing may be subject to human’s biased interpretation. Reliable open-source software to extract and analyze morphometric data with minimal need for manual intervention are of interest to process a batch of neuronal images.

We present here an automated image analysis routine for segmentation, classification and quantification of the morphology of sparsely labelled, cultured neurons based on several novel image processing algorithms. The developed software offers a user-friendly GUI that lets the user interact easily with different tools which are chained together. After segmentation, our software automatically classifies all reconstructed neurites and saves a list of key output parameters to .csv and .txt file formats for consecutive analysis with statistical software. All resulting figures and illustrations are also stored as .tiff files within the defined output folders. The software is fully-automated with minimal need for manual intervention, enabling the analysis of hundreds of neuronal images per day.

For most neuronal data, the built-in segmentation module can fulfil segmentation quality expectations, eliminating the need to run any auxiliary segmentation software or tool before starting the neurite classification step. However, the accuracy achieved in by the proposed neurite classifier depends most strongly on the outcome of the segmentation step. To this end, we included optional pre-processing and gap-bridging modules to reduce the effects of noise and artifacts. For complicated cases such as multiple overlapping cells or very low signal to noise, more powerful and tunable standalone segmentation software may produce more accurate results. In such cases, instead of importing the original raw images, users can load binarized image files, which have been segmented using another software/tool, with the single requirement that the segmentation has to contain a reconstructed soma. Our software then measures total soma area and extracts information on axon, dendrite, axonal and dendritic arbors with a high accuracy. In this way, the proposed approach could be applied as a plugin or extension to open-source frameworks such as Vaa3D or commercial neuron analysis and visualization frameworks.

The classification module robustly detects and measures soma and neurites in 98.5% of tested cases. Furthermore, it accurately reproduces relevant quantitative readouts produced in manual analysis by an expert user and is sensitive enough to detect differences between biological treatments groups. However, it does not distinguish crossing neurites from branches due to lack of three-dimensional information. This overestimates the number, and under-estimates the length of individual branches. The total length of branches is nevertheless detected accurately. One possible solution to this problem, could involve manual post-processing of the classification result. As another solution, one could define rule-based decision-making algorithms based on image features like diameter, signal intensity, curvature, or the propagation vector of the intersecting neurites in order to distinguish and label them individually.

Furthermore, due to the composition of the test dataset of early developing neurons that have elaborated axons but comparably underdeveloped and short dendrites, we were not able to objectively score the performance for mature dendrite quantification. It is possible, that dendrites might be affected more by the crossing versus branching problem described above due to their proximity especially proximal to the soma. Furthermore, the order of classification, with axon branches being classified before dendrite branches, likely skews accuracy towards axon morphology. Users should therefore carefully monitor the performance of this tool when assessing dendrite morphology and, in case phenotypes are not accurately captured, consider more dendrite-specific analysis strategies such as Sholl analysis. It is conceivable, however, that the software can be readily applied to quantifying growth potential, polarization, or protrusion density of neurons at earlier developmental timepoints. When used with a shorter minimal length threshold and subsequent filtering by length it could also measure density of branch-precursor structures such as filopodia in addition to branches individually. In the future the detection of dendritic spine number, size and length could be an avenue to pursue.

The computational complexity of neurite branching analysis scales exponentially with neuron size. The current implementation allows for a high-throughput analysis of early developmental timepoints when neurons are small – without even exhausting the processor and RAM of a standard laptop. The analysis can easily be run in the background while continuing working on the machine. In case larger neurons are of interest, however, the duration might extend considerably, even when using more powerful workstations. We nevertheless believe that the benefits of fully automated implementation outweigh the downsides of analysis times and suggest batch-processing of large datasets overnight in such cases.

In conclusion, we present a modular, fully automated software which provides reliable segmentation, classification, and quantification of the morphology of cultured neurons from two-dimensional fluorescence micrographs. We have evaluated the performance of the proposed neurite segmentation and classification approach by comparing it to that of an expert human user and found a high similarity in reconstruction and extraction quality. The fully automated implementation of the software will facilitate the quantification of large datasets containing micrographs of more neurons from multiple experiments to improve statistical confidence and enable fast screening of multiple treatments and thereby accelerate and improve research on neurodevelopmental mechanisms.

## Software Availability

The software code and example raw images for testing the software are available in the software repository: https://github.com/AG-Ewers/NeuronAnalysis

## Supplemental Materials

### Supplementary Methods

#### Neuron culture, microscopy and pre-processing of the validation dataset (Brosig et al., 2019)

Primary hippocampal neurons of either C57 BL/6 or C57 Bl/6 PRG2 -/- mice were cultured on laminin and poly-ornithine coated coverslips at a density of 15,000 cells/cm^2^. To visualize individual cells, neurons were transfected with a plasmid encoding for green fluorescent protein (GFP) after two days in culture. Neurons were fixed after five days in culture and immunolabelled for the axon marker Tau and GFP. Individual neurons were imaged using a Nikon Eclipse Ti epifluorescence microscope with a 40x objective (NA: 0.8). Prior to analysis, the GFP channel was converted to an 8-bit tif. Manual tracing and classification of neurites was performed using NeuronJ (Meijering et al., 2004), a widely-used ImageJ plugin based on a ridge-finding algorithm which allows semi-automated centerline tracing of 2D neuron images (Narro, 2007). Neurites in the ground truth data were classified as primary axon (Tau-positive and longest process originating from soma), dendrites (other processes originating from soma), primary axon branches (processes longer than 10 µm originating from primary axon), secondary, tertiary, and quaternary branches as well as dendritic branches (originating from dendrites).

## Supplementary Figures

**Figure S1:**
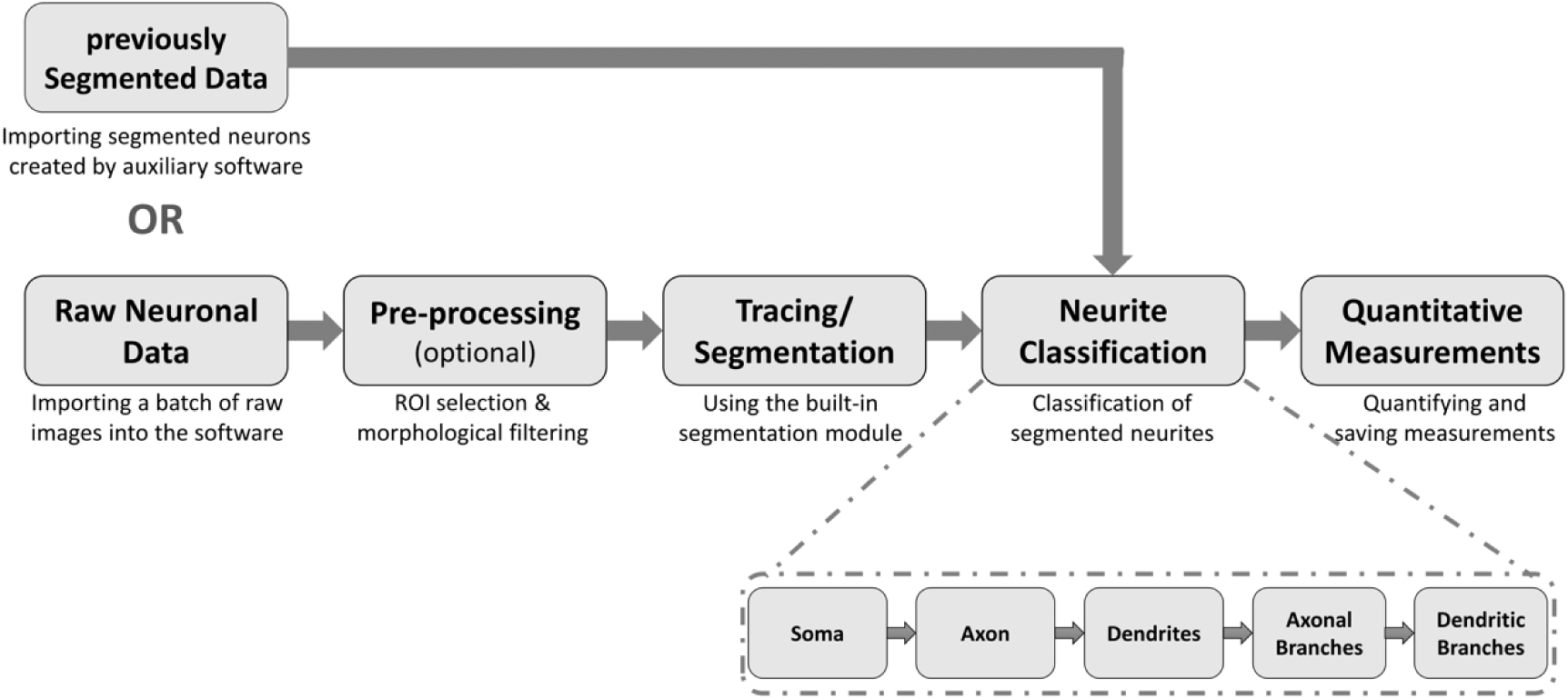
Scheme of the workflow for neuron reconstruction and neurite classification. Shown are the major steps offered by the software and the respective options for the input of raw or preprocessed data and the automated saving of batch processed data.

**Figure S2:**
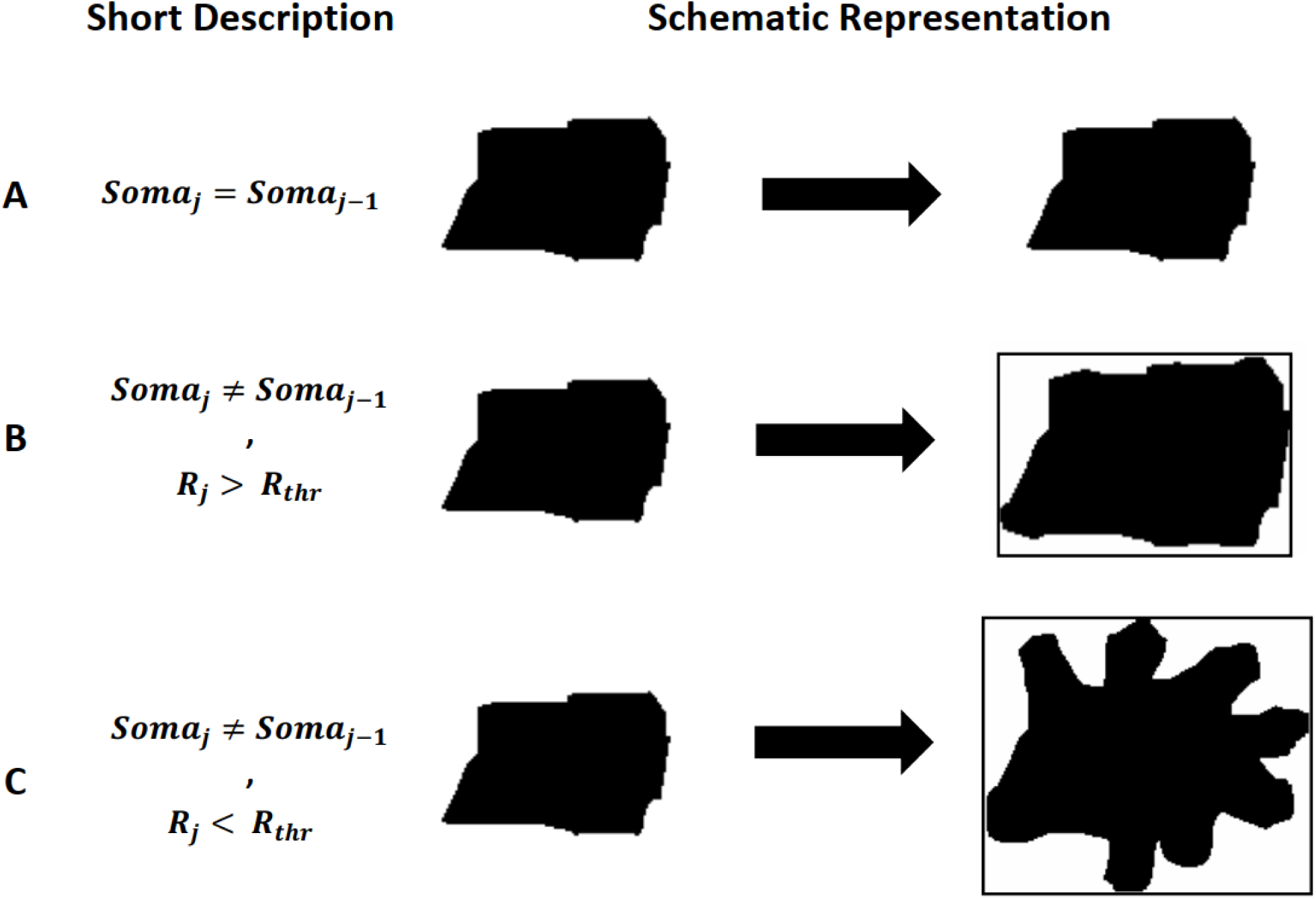
Different approaches for filling the soma. **(A)** Updating the soma does not change its size or shape anymore. This so-called ‘saturated’ case results in breaking the soma expansion process; **(B)** After analyzing the outer vicinity of the current soma, it is now expanded to cover some of the neighboring pixels as well. Since the ratio between the ‘size of the updated soma’ and the ‘size of the bounding box’ (i.e. ***R***_***j***_) is still lower than a pre-defined threshold (i.e. ***R***_***thr***_), the expansion process will continue; **(C)** Similar to the previous case, the reconstructed soma is still expanding. However, the relatively low value of ***R***_***j***_ shows that the soma mask tends to overlap with some non-soma regions, leading to transitions from soma to dendrites and/or the axon. As a result, the soma expansion should stop.

## Notes

### Competing Interest Statement

The authors have declared no competing interest.

